# Excess of mutational jackpot events in growing populations due to gene surfing

**DOI:** 10.1101/053405

**Authors:** Diana Fusco, Matti Gralka, Alex Anderson, Jona Kayser, Oskar Hallatschek

## Abstract

One of the hallmarks of spontaneous mutations in growing populations is the emergence of mutational jackpot events - large mutant clones arising from mutations that by chance occur early in the development of a cellular population so that their progenitors benefit from prolonged growth. Due to their sheer size, these jackpot events, first discovered by Luria and Delbrück [1], are thought to have momentous roles in short-term evolutionary processes, including the adaptation from standing variation [2–4], evolutionary rescue [5], drug resistance evolution [6–10], and the somatic evolution of genetic diseases [11, 12]. However, because the emergence of jackpot events has been understood only in uniformly growing populations [1, 10, 13], it is currently impossible to predict their impact on the evolution of many naturally structured populations. To study jackpot events in spatially structured populations, we tracked mutant clones in microbial populations using fluorescent microscopy and population sequencing. High-frequency mutations were massively enriched in microbial colonies compared to well-shaken liquid cultures, as a result of late-occurring mutations surfing at the edge of range expansions [14–16]. We provide a mathematical theory that explains the observed excess of jackpot events and predicts their role in promoting rare evolutionary outcomes. In particular, we show that resistant clones generated by surfing can become unleashed under high selection pressures, and thus represent a drug resistance hazard for high-dose drug treatments. An excess of mutational jackpot events is shown to be a general consequence of non-uniform growth and, therefore, could be relevant to the mutational load of developing biofilm communities, solid tumors and multi-cellular organisms.

---

In the original Luria-Delbrück experiment, the size of mutational jackpot events was measured by counting single colonies on selective plates. As a simpler mean towards the same end, we employed population sequencing with low error rates (Appendix B), which returns frequencies of new mutations on many genomic sites simultaneously and independently of their phenotypic effect. Specifically, we sequenced populations of a mutator strain of *E. coli* cultured in well-mixed liquid, where growth is uniform, and on solid agar medium, where most growth occurs at the colony edge (Appendix B) [17, 18]. By growing from a small number of initial cells to a similar final size, all populations went through a comparable number of cell divisions in the range of several billion divisions (Supplementary Table B2). By counting the observed frequencies of Single Nucleotide Polymorphisms (SNPs) in the populations, we obtain the number of sites in the genomes where the clonal sub-population carrying the derived mutation has a frequency larger than a given frequency value *x*, shown in (Fig. 1b. Our deep sequencing procedure allows us to detect all clones that have frequencies larger than about 10^−3^, yielding at least 600 such high-frequency events in each colony, which characterize the statistics of jackpot events.

**Figure 1.**
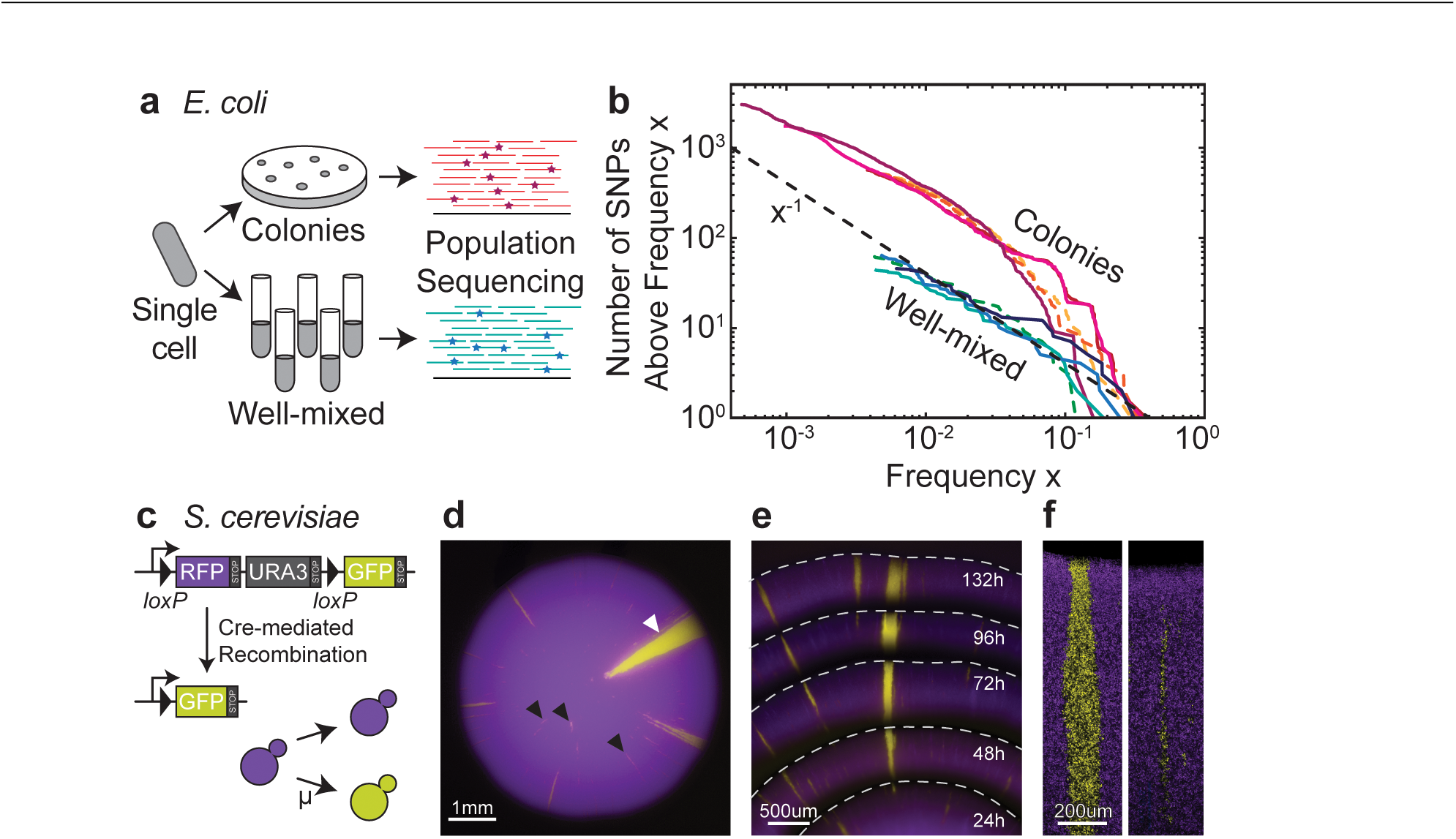
Gene surfing promotes the occurrence of mutational jackpot events in colonies. **(a)** Starting from few cells of a mutator strain of the bacterium *E. coli* (*mutT* deletion, Appendix B), we grew 4 colonies and 4 liquid cultures up to an average population size of 3 × 10^9^ cells (Supplementary Table B2). We sequenced the DNA from each of these populations at a coverage of at least 1000× and measured the frequency of single nucleotide polymorphisms (SNPs) that arose during the growth process. The number of SNPs that occur at a frequency higher than *x* is displayed in panel **(b)** (colonies in warm colors, well-mixed in cold colors), which shows that colonies produce an excess of high-frequency mutations: We found about 10 times more mutants above a frequency of 1% in colonies than in liquid culture (dashed green vs. dashed light orange). Colonies twice and three times larger (solid magenta and purple lines, respectively) similarly exceed the well-mixed null expectation (dashed black line, fit to the well-mixed data with a mutation rate of *μ* = 0.4 per genome per replication). (**c-f**) An engineered strain of S. *cerevisiae* that stochastically switches from RFP (purple) to GFP (yellow) at a rate of 10^−3^ per cell division, enables us to visualize high frequency jackpot events as they arise during colony growth. Monitoring the spatial distribution of mutants using fluorescence time-lapse microscopy (panel e, see also Supplementary Movie 1) reveals that mutant clones come either as sectors with actively growing front regions (**f**, left; **d**, white arrow) or as “bubbles” (**f**, right; **d**, black arrows), which are non-growing mutant clones that have lost contact to the range edge.

Because mutation rates did not significantly vary between the two modes of growth (liquid vs. colony, Supplementary Table B1), one expects the same total number of mutational events in populations grown to the same final size. Yet, (Fig. 1 shows that colonies 1 and 2 have approximately ten times more mutant clones above frequencies of 1% - corresponding to clones of at least 10^7^ cells - compared to the equally large well-mixed population. We consistently find a similar excess of high-frequency clones across samples even if the population size varies.

To reveal the nature of the high-frequency clones in colonies, we monitored the spatial distribution of mutant clones using separate fluorescent marker experiments. We employed a genetically engineered budding yeast strain capable of switching from a red-fluorescing state to a green-fluorescing state at a rate of ≈ 1.6 x 10^−3^ per replication [19]. This heritable, nonreversible switch is mediated by the stochastic expression of Cre recombinase (see (Fig. 1c and Appendix B for details). As shown in (Fig. 1, the resulting colonies exhibited both elongated speckles, which we term “bubbles”, as well as previously described spoke-like sectors [17]. Importantly, image analysis of the clone area obtained from 343 colonies yields a histogram that is consistent with the shoulder-like distribution obtained from our sequencing approach (Supplementary Fig. B3). Thus, both the fluorescent data from just one “engineered” site and the sequencing data covering many genomic sites seem to reflect the same mechanism shaping the clone size distribution.

The fluorescence data, moreover, reveals where clones emerge and how they grow. Time-lapse movies show that most high-frequency clones first arise near the front of the growing colony ((Fig. 1e, Supplementary Movie 1). The resulting clonal patches grow with the advancing frontier until they lose contact to the advancing frontier, upon which they become trapped as bubbles in the non-growing bulk of the population. Rarely, clones are able to “surf at the front until the end of the experiment and give rise to sectors. Such allele surfing is a characteristic feature of range expansions [14, 16, 20] and has been demonstrated to be pervasive in microbial communities [17, 21–23].

To understand how gene surfing generates clones of different sizes, we studied their emergence in two-dimensional range expansion simulations (three-dimensional results are provided in Appendix A). Specifically, colony growth is implemented by the random addition of new demes to the advancing frontier. The newly added deems inherit their ancestral genotype unless they mutate, which occurs at a fixed rate (Appendix B; Supplementary Movie 2). Interestingly, this simple meta-population model gives rise to a clone size distribution with two distinct power-law regimes that correspond to bubbles and sectors, respectively, as shown in (Fig. 2. We derive the precise value of the two power-law exponents in Appendix A from the characteristic fractal properties of sector boundaries [24], which have been previously measured [17]. Our scaling arguments predict that the clone size distributions obtained for different population sizes and mutation rates should collapse onto one master curve when the clones frequencies are measured in terms of the characteristic frequency of the largest occurring bubble *x_c_*. Indeed, after rescaling, our sequencing data show remarkable agreement with the scaling function obtained from our simulations ((Fig. 2).

**Figure 2.**
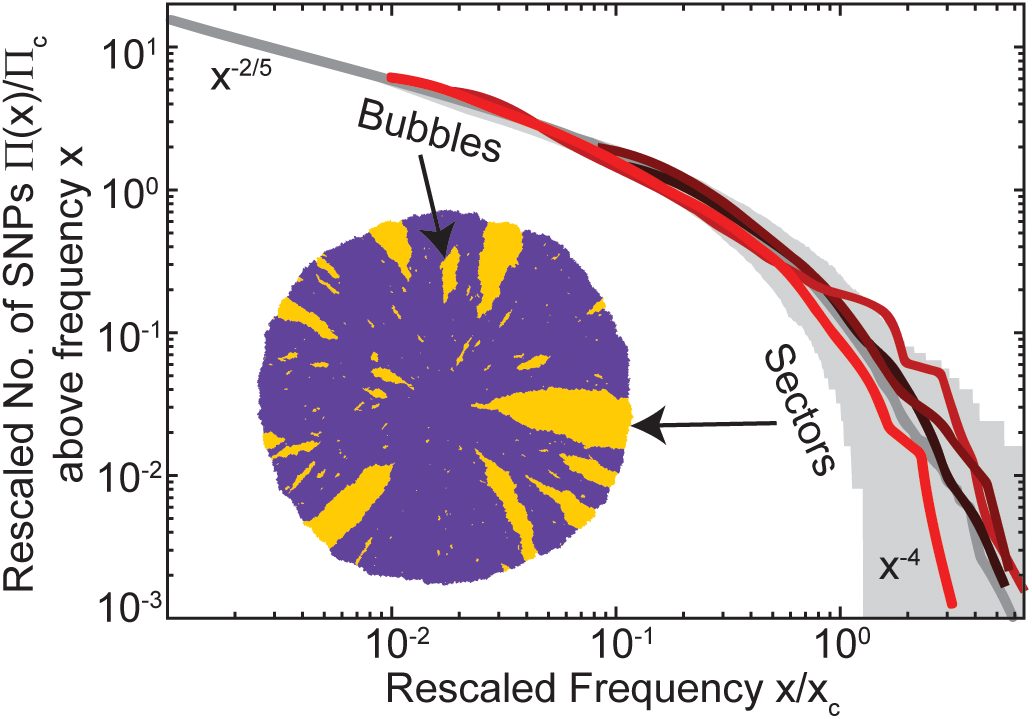
Gene surfing theory explains the excess of jackpot events and predicts evolutionary consequences. In our coarse-grained simulations, new mutations only arise and surf at the edge of the colony. Depending on the duration of surfing, mutants generate spatially encased bubbles or continually growing sectors. The simulated mutant histogram (gray solid line) exhibits two predictable power-law tails that characterize bubbles and sectors, respectively (Appendix A). The experimental mutant spectra (colored lines) nearly collapse onto the theory line upon scaling the two axes by the frequency of the largest bubble *x_c_* and the probability Π_*c*_ of observing a clone larger than *x_c_*, respectively (shaded area represent the 95% confidence interval, see Appendix B for details).

The primary consequence of more jackpot events is that, typically, the total number of mutants will be much larger in a population of given size. This leads to a greater probability for rare evolutionary outcomes as summarized in Box *Theory* and Appendix A. Moreover, evolutionary dynamics is influenced by mutational jackpot events not only because of their size but also because of their particular spatial structure. The emergence of sectors, which sporadically arise from neutral mutations, is strongly suppressed when mutants carry a cost (Supplementary Fig. A3), commonly observed for drug resistance in the absence of antibiotics [26]: Deleterious mutants can surf only briefly before they are overtaken by faster-growing wild-type cells and fall behind the growing frontier.

In the absence of antibiotics, then, costly resistant clones are expected to reside in bubbles, encased by an expanding wild-type population. Upon a sudden environmental change, however, e.g., by a strong antibiotic attack killing the susceptible wild-type, the trapped mutants may become unleashed, regrowing and thus rescuing the population from extinction. At intermediate drug concentrations, sufficiently strong to slow down proliferation of the wild-type without eradicating it, we expect minimal net population growth and successful containment of resistant cells. Indeed, when we implement drug treatment as an increased death rate for wild-type cells ((Fig. 4a, Supplementary Movie 3), our simulations show the smallest net population growth for intermediate death rates, while the populations grow to large sizes for both low and high death rates.

**Figure 4.**
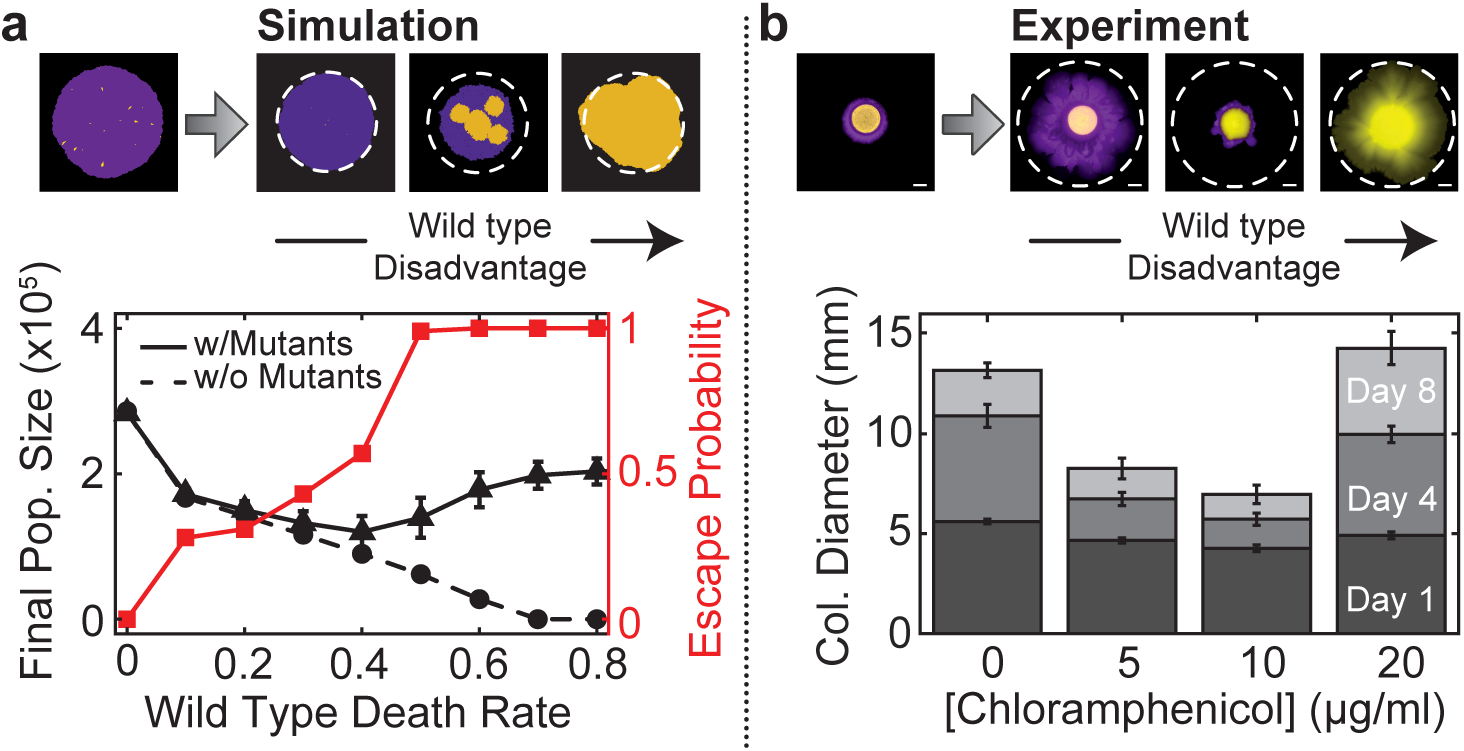
Simulations suggest that high drug concentration treatments could promote the spreading of resistance by unleashing mutants that are otherwise trapped within the colony bulk when the mutants carry a cost in the absence of a selection pressure, most mutants in our simulations are trapped in non-growing bubbles rather than sectors. Starting with pre-grown colonies (black solid triangles) and resistance-free control colonies (black dashed circles) as initial condition, we simulated the effects of treatment with antibiotics of varying concentrations by tuning the death rate δ of the wild type (Appendix B). After 140 generations, resistance-free colonies showed decreasing population size up to complete eradication for δ > 0.7, as expected. Conversely, pre-grown colonies that contained resistant mutants within the bulk, exhibited an initial decrease in population size followed by a steady increase for δ > 0.4 (error bars represent standard deviations over 100 simulations). Visualization of the simulated colonies shows that in this range of death rates, the previously encapsulated mutants escape the surrounding wild type (red squares) and can then grow indefinitely (Supplementary Movie 3). A high dose of antibiotics may thus not only fail to eradicate the population, but even promote the spreading of resistance. This effect can be replicated in simple conceptual experiments with *E. coli*, shown in (b), which demonstrate that high antibiotic concentrations can unleash trapped mutant clones. A droplet of resistant cells (yellow) embedded in a larger droplet of susceptible cells (purple) was inoculated at different antibiotic concentrations. After 8 days of growth intermediate antibiotic concentrations exhibited the least amount of total population growth. The highest drug concentrations eradicated the wild type and thus allowed the resistant mutants to spread freely. Error bars represent standard deviations over 16 replicates and scale bars correspond to 2 mm.

The hypothesis that growth control may sometimes be more effective than complete eradication has been recently proposed in the context of cancer [27, 28] based on mathematical modeling of exponentially growing tumors when treated with a varying dose of chemotherapy over time. Our simulations provide a uniquely spatial mechanism of how excessively high drug concentrations can promote the spreading of drug resistance. This effect can be demonstrated in conceptual experiments in which resistant cells are embedded within a colony of susceptible cells. We find that the resistant cells stay trapped even at intermediate drug concentrations that severely limit the wild-type growth. For higher drug concentrations, the mutants are released and rapidly grow ((Fig. 4b). Hence, to optimally curtail microbial growth in our particular setup, the drug concentration should indeed be set at an intermediate sweet spot.

In combination with a generalization of the Luria-Delbrück theory, our experiments suggest that spatially growing populations exhibit an excess of high-frequency clones due to allele surfing. These high-frequency clones come as growing sectors and non-growing bubbles. Bubbles are spatially encased by wild-type cells and represent a concealed drug resistance hazard as they can become unleashed under high drug concentrations. Our theory suggests that the excess of jackpot events is not limited to microbial colonies but is expected to arise generally in populations that exhibit non-uniform growth rates.

In addition to antibiotic resistance evolution in pathogenic biofilms, an excess of jackpot events could thus also be relevant during the somatic evolution of some types of cancer [12, 29], as growing solid tumors often exhibit less growth in necrotic core regions [30] and sectoring has been recently documented [31] (for the three-dimensional predictions of our jackpot theory see Appendix A). Our results show that deviation from the classical Luria-Delbrück theory may simply indicate non-uniform growth rather than non-neutral evolution, in contrast to what has been recently hypothesized in the context of tumor evolution [12].

**Origin and statistical consequences of the excess of jackpot events**

That range expansions generate a larger mean number of mutants can be understood from a simple mathematical argument. If mutations arise at a low rate *μ* per cell division as the population is growing to a final size N, one expects roughly *μ*N mutational events to occur. In each generation, the frequency of mutants in the population increases by *μ*, on average. In a well-mixed, uniformly grown population, the number of generation is log_2_ N, and hence the expected total number of mutants in this case is proportional to *μ*N log_2_ N. In a range expansion, only cells near the edge of the colony have access to sufficient nutrients, leading to the formation of a layer of growing cells of width λ (in units of cell diameters). Since a length λ is added per generation to the radius of the colony, we can estimate that *R*/λ ∝ N^1/2^/λ generations elapse at the frontier during the growth process. Hence, the final total number of mutants created during a range expansion is proportional to *μ*N^3/2^/λ, which for large N is much larger than in a uniformly grown population of the same size.

However, as in the classical Luria-Delbrück case, the mean is usually not a useful quantity because it is dominated by very rare, large events. Nevertheless, both the typical and the mean number of mutations exceed the well-mixed expectations, as our stochastic analysis shows ((Fig. 3 and Appendix A).

Complex phenotypes, such as drug resistance or the onset of cancer, often require the accumulation of multiple mutations, for which jackpot events can be key [11, 25]. Range expansions may be favorable for the acquisition of secondary mutations because the pool of individuals carrying the first (driver) mutation is, both on average and typically, larger than in uniformly grown mutation ((Fig. 3a). As shown in (Fig. 3b, the probability of secondary mutations can be almost an order of magnitude larger in spatial population compared to well-mixed ones, especially when mutation rates are low.

**Figure 3.**
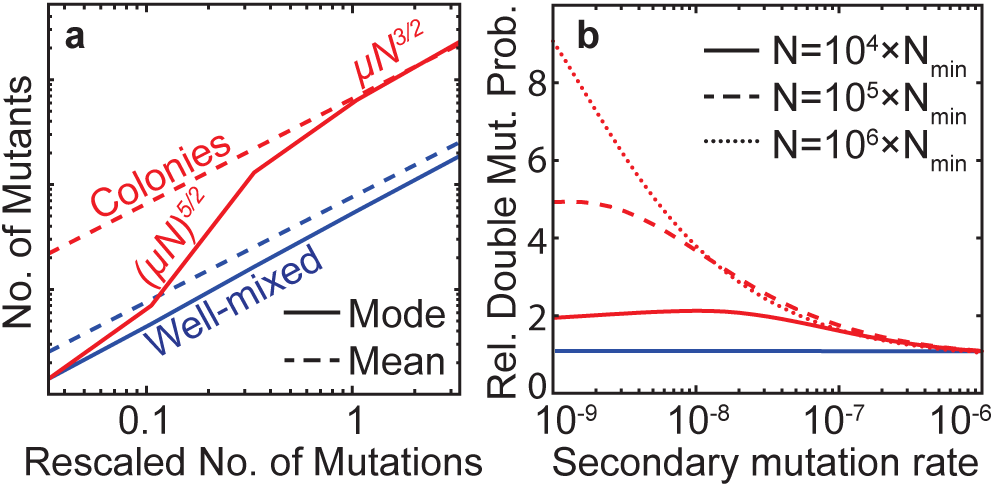
Gene surfing theory predicts an excess of jackpot events and its consequences. **(a)** By sampling the distribution of clone sizes Π obtained for the sequenced colonies (Appendix B), we compute the typical (mode, solid line) and the mean (dashed line) number of mutants as a function of the rescaled number of mutations *μN*Π_*c*_. We find that the typical number of mutants scales much faster in colonies (red) compared to liquid cultures (blue), approaching the average number of mutants when the probability of sampling at least one sectors is *O*(1) (*μN*Π_*c*_ ≈ 1). Therefore, given the same number of mutations, range expansions carry many more mutants than equally large uniformly grown populations. **(b)** The excess of large-frequency clones promotes multi-step evolutionary process, such as the emergence of double mutants. We find that the relative probability (compared to the well-mixed expectation) of producing a double mutant is always higher in a colony, especially when the second mutation rate is low. Here, *N*_min_ = *x*_min_*N* represents the minimum population size below which fine scale effects play a role and our theory cannot be directly applied (Appendix A). In our *E. coli* sequenced colonies, we find *N*_min_ ≈ 10^5^ (Supplementary Table B3 and Supplementary Fig. B5).

